# Production of novel biosurfactant by a new yeast species isolated from *Prunus mume* Sieb. *et* Zucc

**DOI:** 10.1101/2022.03.15.484497

**Authors:** Jeong-Seon Kim, Miran Lee, Dae-Won Ki, Soon-Wo Kwon, Young-Joon Ko, Jong-Shik Kim, Bong-Sik Yun, Soo-Jin Kim

## Abstract

Biosurfactants reduce surface and interfacial tension due to their amphiphilic properties, and are an eco-friendly alternative for chemical surfactants. In this study, a novel yeast strain JAF-11 that produces biosurfactant was selected using drop collapse method, and the properties of the material were investigated. The nucleotide sequences of the strain were compared with closely related strains and identified based on the D1/D2 domain of the large-subunit rDNA (LSU) and internal transcribed spacer (ITS) regions. *Neodothiora populina* CPC 39399^T^, the closest species with strain JAF-11 in the phylogenetic tree, showed a sequence similarity of 97.75% for LSU and 94.27% for ITS, respectively. The result suggests that the strain JAF-11 represent a distinct species which cannot be assigned to any existing genus or species in the family *Dothideaceae*. Strain JAF-11 was able to produce biosurfactant reducing the surface tension of medium to 34.5 mN/m on the 6^th^ day of culture and the result of measuring the critical micelle concentration (CMC) by extracting the crude biosurfactant was found to be 24 mg l^-1^. The molecular weight 502 of the purified biosurfactant was confirmed by measuring the fast atom bombardment mass spectrum (FAB-MS). The chemical structure was analyzed by measuring ^1^H nuclear magnetic resonance (NMR), ^13^C NMR, two-dimensional NMRs of the compound. The molecular formula was C_26_H_46_O_9_, and it was composed of one octanoyl group and two hexanoyl group to myo-inositol moiety. The new biosurfactant is the first report of a compound produced by a novel yeast strain JAF-11. This new biosurfactant is proposed as potential candidate for use in a variety field.

## Introduction

The surfactants have amphiphilic properties possessing both non-polar (hydrophobic) and polar (hydrophilic) moieties that allows reducing the surface and interfacial tension between biphasic systems as liquid-liquid interface or solid-liquid boundaries [1, 2]. The surfactants are one of the important compounds having potential commercial application of detergents, cosmetics, food ingredients, agricultural, pharmaceutical, paint, textile and paper etc. [3, 4]. However, surfactants are mainly chemically synthesized from petroleum-based resources, which can cause environmental problems due to toxicity [5]. As environmental awareness has gradually increased over the past decades, the demand for eco-friendly compounds has increased, and accordingly, there is an increasing interest in microbial biosurfactants [6]. In particular, microbe-derived biosurfactants have advantages such as environmental compatibility, activity, stability, and lower toxicity compared to chemically synthesized equivalents [7–11].

Biosurfactants are classified according to ionic charges (anionic, cationic, non-ionic and neutral biosurfactants), molecular weight (high molecular and low molecular weight biosurfactants), secretion type (intracellular, extracellular and adhered to microbial cells), and chemical structure (glycolipids, lipopeptides, fatty acids, phospholipids, neutral lipids and polymeric biosurfactants), and it has also been previously reported that various biosurfactants are produced depending on the type of microorganisms [12]. Among them, glycolipids have been best known as a class of biosurfactants that exhibit the highest commercial potential due to the high microbial productivity [13]. The microbe-derived biosurfactant rhamnolipids, also known as glycolipids, was described for the first time in 1946 [14]. Despite their high productivity, most of them is produced by opportunistic pathogen *Pseudomonas aeruginosa* [15]. The sophorolipids belonging to the group of glycolipid are one of the most promising biosurfactant because they are produced by non-pathogen yeast *Starmerella bombicola* and other *Candida* spp. (*Candida stellate, C. riodocensis, C. apicola, C. batistae, C. kuoi,* and *C. floricola*) [16, 17]. Mannosylerythritol lipids (MEL) by a variety of *Pseudozyma* yeasts and trehalolipids by *Rhodococcus erythopolis* have been reported as representative biosurfactants belonging to the glycolipid [18]. It has been reported that microorganisms such as the genera *Acinetobacter*, *Arthrobacter*, *Pseudomonas*, *Halomonas*, *Bacillus*, *Rhodococcus*, *Enterobacter*, and few yeast genera produce not only glycolipids but also other types of biosurfactants [19].

In the 20^th^ century, research on biosurfactants focused on understanding and optimizing the production process, and since then, research on utilizing various renewable resources, discovering new producer strains, or developing genetically modified strains has been reported [6]. Since biosurfactants have a wide range of functional properties depending on their chemical structure [19], it is necessary to continuously secure a variety of biosurfactants in order to be applied to various fields. The present study is the first report of novel biosurfactant extracted from the novel yeast strain JAF-11. We isolated a new species of yeast that produces biosurfactant from flower in order to secure various biosurfactants, and chemical structure of the biosurfactant extracted from the yeast has been characterized and identified as a novel biosurfactant by nuclear magnetic resonance spectrometry techniques.

## Materials and methods

### Culture medium for biosurfactant production by yeast

The culture medium composition used in these studies for biosurfactant-producing yeast was as follows (w/v): glucose (1.5%), soybean oil (1.5%), ammonium sulfate (0.1%), potassium phosphate (0.25%), sodium phosphate (0.01%), magnesium sulfate (0.05%), calcium chloride (0.01%), manganese sulfate (0.002%) and peptone (0.1%).

### Isolation and screening of biosurfactant producing yeast

The biosurfactant-producing yeast used in this study was isolated from flower (*Prunus mume* Sieb. *et* Zucc.) collected from apricot village in Gwangyang, Republic of Korea during March 2018. The biosurfactant was screened using modified drop collapse method as follows: 100μL of the culture supernatant and water (1:1, v/v) was pipetted and placed on parafilm [20, 21]. The selected strain JAF-11 was maintained for storage at −80°C in 15% (v/v) glycerol, and was deposited with the patent depository as KACC 83047BP.

### Identification of biosurfactant producing yeast

The identification of strain JAF-11 was conducted based on multigene phylogenetic analysis of the nucleotide sequences combined with the D1/D2 region of large-subunit (LSU) and the internal transcribed spacer (ITS) region ribosomal DNA (rDNA) genes. The DNA sequencing was performed by Macrogen Inc. (Seoul, Korea) and the gene sequences of related species were retrieved from GenBank database. The phylogenetic tree was inferred by using the maximum likelihood method with 1,000 bootstrap replicates and sequences analysis was performed in MEGA X software.

### Time course of the growth and determination of surface tension

The strain JAF-11 was grown in 500 ml culture medium at 25°C on a rotary shaker at 150 rpm, and then the optical density at 600 nm was measured for 8 days by a UV spectrophotometer. To confirm the production of biosurfactant, the culture medium of strain JAF-11 was prepared through a 0.22 *μ*m filter and measured the surface tension(ST) every day. For ST measurement, the Wilhelmy plate method was used at room temperature with a force tensiometer K11 (Krüss, Germany). All tests were performed in triplicates.

### Measurement of critical micelle concentration (CMC) of biosurfactant

To extract the crude biosurfactant, 8 L culture medium inoculated with freshly grown strain JAF-11 was mixed with HP-20 non-polar resin (Mitsubishi chemical, Japan) and eluted in methanol. After removing methanol by rotary vacuum evaporation, the concentrated solution was partitioned with an equal volume of ethyl acetate. The ethyl acetate was concentrated under vacuum evaporator, and the product was used for CMC measurement after purification using Flash silica gel column chromatography (CHCl_3_:MeOH, 50:1 to 1:1, v/v, stepwise) (SK chemical, Korea). The crude biosurfactant were dissolved in distilled water and serially diluted to concentrations in the range of 0-250 mg l^-1^. The CMC was determined by plotting the surface tension against the log of the biosurfactant concentration using a force tensiometer K11 (Krüss, Germany) [22].

### Isolation and purification of biosurfactant compound

A yeast strain of JAF-11 was cultured in a medium described above for 5 days at 25°C on a rotary shaker incubator at 150 rpm. The culture broth was subjected to Diaion HP-20 column chromatography and eluted with 30% aq. MeOH, 70% aq. MeOH, MeOH, and acetone. The biosurfactant activity of eluates were evaluated by the drop collapse method, and the MeOH eluate was chosen and concentrated by rotary vacuum evaporator. The concentrate was partitioned between ethyl acetate and water. The ethyl acetate portion showing biosurfactant activity was concentrated and applied to silica gel column chromatography (Merck, Germany) eluted with CHCl_3_:MeOH (50:1, 20:1, 10:1, 5:1, 2:1, and 1:1, v/v) (Fig 1). An active fraction, CHCl_3_:MeOH (20:1), was subjected to Sephadex LH-20 (GE Healthcare, Sweden) column chromatography eluted with CHCl3:MeOH (1:1, v/v). The fractions showing biosurfactant activity were combined and concentrated (Fig 1). The concentrate was further separated by the medium pressure liquid chromatography (MPLC, Teledyne ISCO, USA) equipped with Redisep Rf C_18_ reversed-phase column (Teledyne Isco, USA) using a gradient of 70%→100% aq. MeOH to yield compound **1** (13.4 mg).

**Fig 1.**
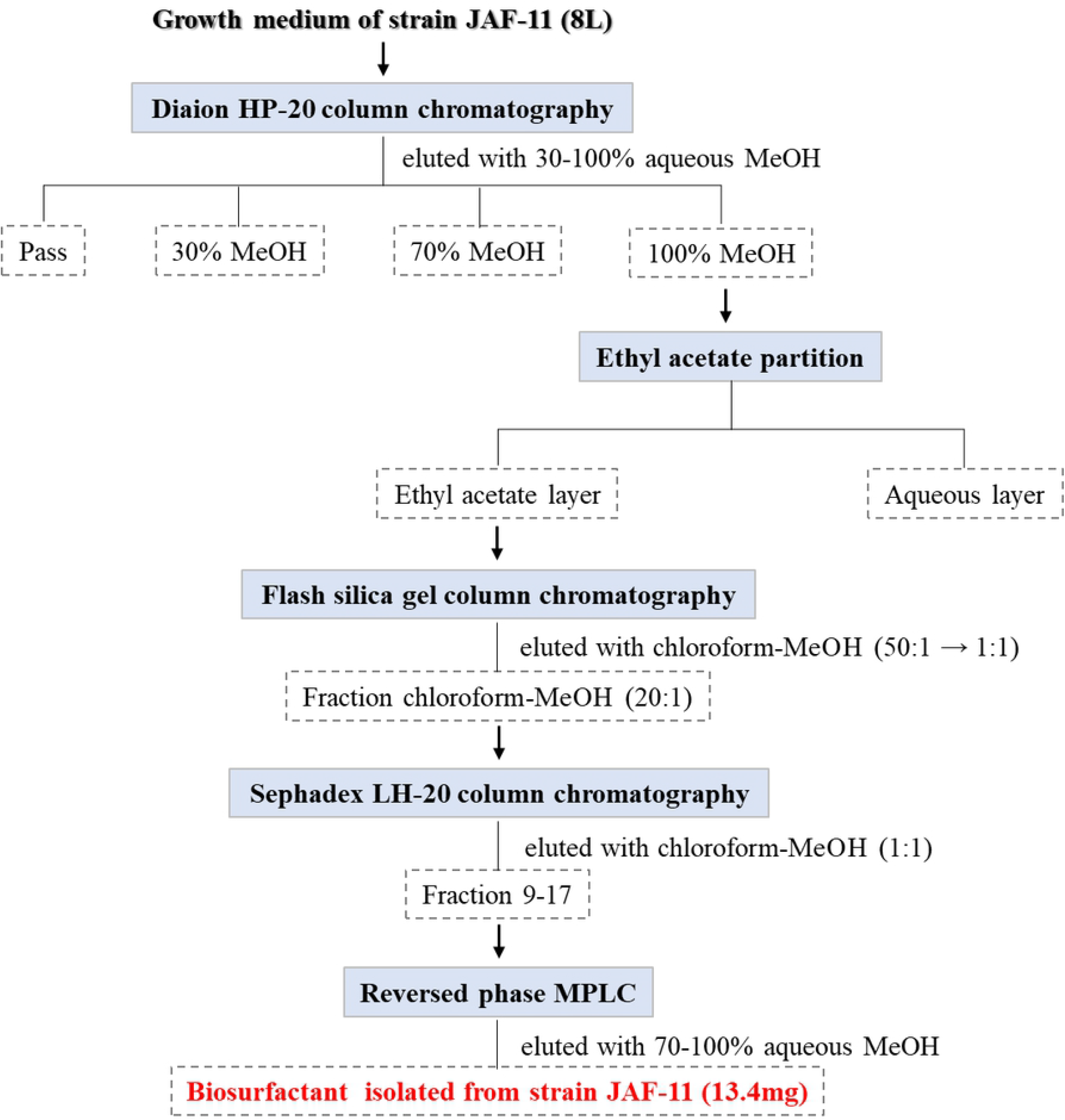
Purification scheme of the biosurfactant produced by strain JAF-11.

### Chemical structure analysis of biosurfactant

The fast atom bombardment mass spectrum (FAB-MS) and high-resolution fast atom bombardment mass spectrum (HRFAB-MS) for the molecular weight and molecular formula, respectively, were measured using a JMS-700 MStation (JEOL, Japan) mass spectrometry. The nuclear magnetic resonance (NMR) spectra were obtained on a JEOL JNM-ECZ500R, 500 MHz FT-NMR spectrometer at 500 MHz for ^1^H NMR and at 125 MHz for ^13^C NMR in CD_3_OD (Andover, USA). Chemical shifts are given in ppm (δ) with tetramethylsilane as the internal standard. For NMR spectra, two-dimensional NMR such as ^1^H-^1^H correlated spectroscopy (^1^H-^1^H COSY), heteronuclear multiple quantum correlation (HMQC), and heteronuclear multiple bond correlation (HMBC) as well as one-dimensional NMR such as ^1^H NMR and ^13^C NMR were employed.

## Result and discussion

### Isolation and identification of biosurfactant producing yeast

Strain JAF-11 was isolated from *Prunus mume* Sieb. *et* Zucc. in Republic of Korea and was selected as a potential biosurfactant producer using modified drop collapse method (S1 Fig). Phylogenetic relationships of strain JAF-11 and the closely related strains were inferred using concatenated LSU and ITS sequences [23, 24]. The LSU and ITS sequences from the reference strains were obtained from GenBank database (Table 1). Phylogenetic analysis revealed that JAF-11 belonged to the family *Dothideales* clade, and formed one compact cluster with *Neodothiora populina* CPC 39399^T^, *Rhizosphaera macrospora* CBS 208.79^T^ and *Phaeocryptopus nudus* CBS 268.37 (Fig 2). The LSU region sequence of strain JAF-11 showed the highest similarity with those of *Rhizosphaera macrospora* CBS 208.79^T^ (97.26%; 15nt substitutions in 548 nt) and *Neodothiora populina* CPC 39399^T^ (97.75%; 13nt substitutions in 578 nt). In the ITS region sequences, strain JAF-11 had sequence similarity of 95.15% (26 nt substitutions in 536 nt) with *Rhizosphaera macrospora* CBS 208.79^T^ and 94.27% (31 nt substitutions in 541 nt) with *Neodothiora populina* CPC 39399^T^. According to Duong Vu et al. (2016), the taxonomic threshold predicted to discriminate yeast species is 98.41% in the LSU region and 99.51% in the ITS region [25]. Consequently, we propose that strain JAF-11 represents a novel yeast species of a new genus based on phylogenetic analysis and the taxonomic thresholds of gene sequence identities.

**Table 1.** GenBank accession numbers of species used in phylogenetic analyses

| Species | Strain no. | GenBank accession no. |  |
| --- | --- | --- | --- |
|  |  | LSU | ITS |
| Family Dothideaceae |  |  |  |
| *Coleophoma crateriformis* | CBS 473.69 | MH871117.1 | MH859358.1 |
| *Coleophoma oleae* | CBS 615.72 | MH872293.1 | KU728511.1 |
| *Cylindroseptoria ceratoniae* | CBS 477.69<sup>T</sup> | KF251655.1 | KF251151.1 |
| *Delphinella strobiligena* | CBS 735.71 | MH872074.1 | MH860318.1 |
| *Dothidea insculpta* | CBS 189.58 | MH869284.1 | AF027764.1 |
| *Dothidea sambuci* | AFTOL-ID 274<sup>T</sup> | AY544681.1 | DQ491505.1 |
| *Dothidea sambuci* | DAOM 231303<sup>T</sup> | NG_027611.1 | AY883094.1 |
| *Dothiora cannabinae* | CBS 737.71<sup>T</sup> | MH872076.1 | MH860320.1 |
| *Dothiora corymbiae* | CBS 145060<sup>T</sup> | MK047482.1 | MK047431.1 |
| *Dothiora elliptica* | CBS 736.71 | MH872075.1 | KU728502.1 |
| *Endoconidioma populi* | UAMH 10902 | HM185488.1 | HM185487.1 |
| *Neocylindroseptoria pistaciae* | CBS 471.69<sup>T</sup> | MH871115.1 | MH859357.1 |
| *Neodothiora populina* | CPC 39399<sup>T</sup> | MW175405.1 | MW175365.1 |
| *Phaeocryptopus nudus* | CBS 268.37 | GU301856.1 | EU700371.1 |
| *Plowrightia ribesia* | MFLU 14-0040 | KM388552.1 | KM388544.1 |
| *Pringsheimia smilacis* | CBS 873.71 | FJ150970.1 | MH860390.1 |
| *Rhizosphaera macrospora* | CBS 208.79<sup>T</sup> | MH872971.1 | MH861202.1 |
| *Stylodothis puccinioides* | CBS 193.58 | MH869286.1 | MH857753.1 |
| *Sydowia polyspora* | CBS 116.29 | MH866487.1 | MH855019.1 |
| **Family Aureobasidiaceae** |  |  |  |
| *Aureobasidium leucospermi* | CPC 15180 | JN712555.1 | JN712489.1 |
| *Aureobasidium leucospermi* | CBS 130593<sup>T</sup> | MH877257.1 | KT693727.1 |
| *Aureobasidium proteae* | CPC 2825 | JN712558.1 | JN712492.1 |
| *Aureobasidium pullulans* | MFLUCC 14-0288 | KM461701.1 | KM388542.1 |
| *Aureobasidium pullulans* | CBS 584.75<sup>T</sup> | DQ470956.1 | FJ150906.1 |
| *Kabatiella lini* | CBS 125.21<sup>T</sup> | FJ150946.1 | FJ150897.1 |
| **Family Cladosporiaceae** |  |  |  |
| *Rachicladosporium cboliae* | CBS 125424<sup>T</sup> | MH875168.1 | MH863703.1 |

**Fig 2.**
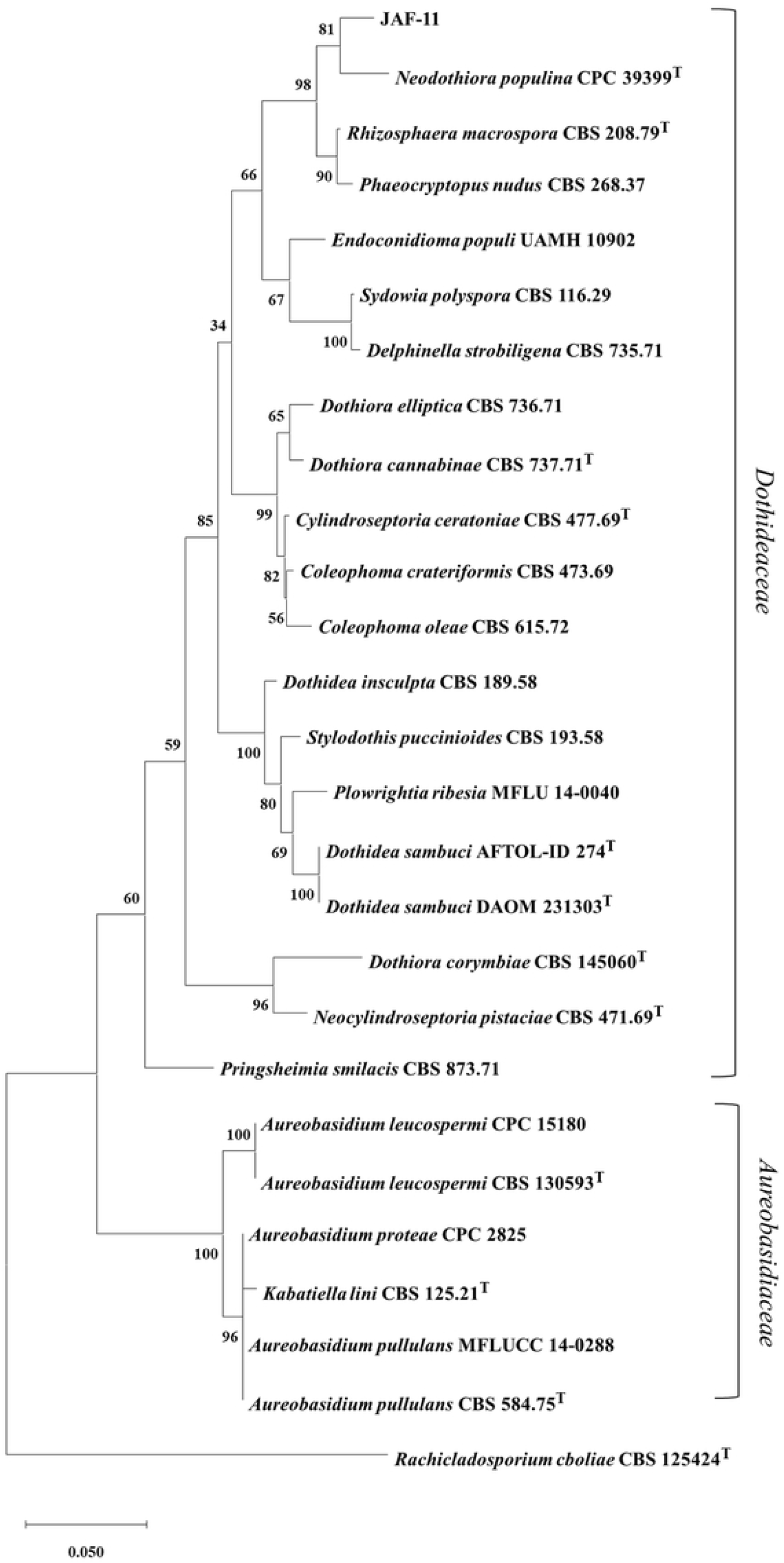
Phylogenetic tree of concatenated LSU and ITS region sequences of closely related species. *Rachicladosporium cboliae* was used as the outgroup in the phylogenetic tree. The phylogenetic tree was constructed using the maximum likelihood method and Tamura-Nei model with bootstrap values 1,000 replicates. The scale bar indicates substitutions per nucleotide position.

### Surface-active properties of biosurfactant

Growth of strain JAF-11 in culture medium was detected by measuring absorbance at 660 nm and surface tensions of the aqueous supernatant was measured using force tensiometer K11 (Krüss, Germany). While the growth of the strain increased continuously for 8 days, the surface tension of the supernatant decreased from 53 mN/m to 34.5 mN/m for 6 days and increased again after 7days. The surface tension recorded the lowest values of 34.5~34.6 mN/m after 5-6 days of incubation (Fig 3). The above results indicated that the highest biosurfactant production in strain JAF-11 is reached at 5-6 days before the stationary growth phase.

**Fig 3.**
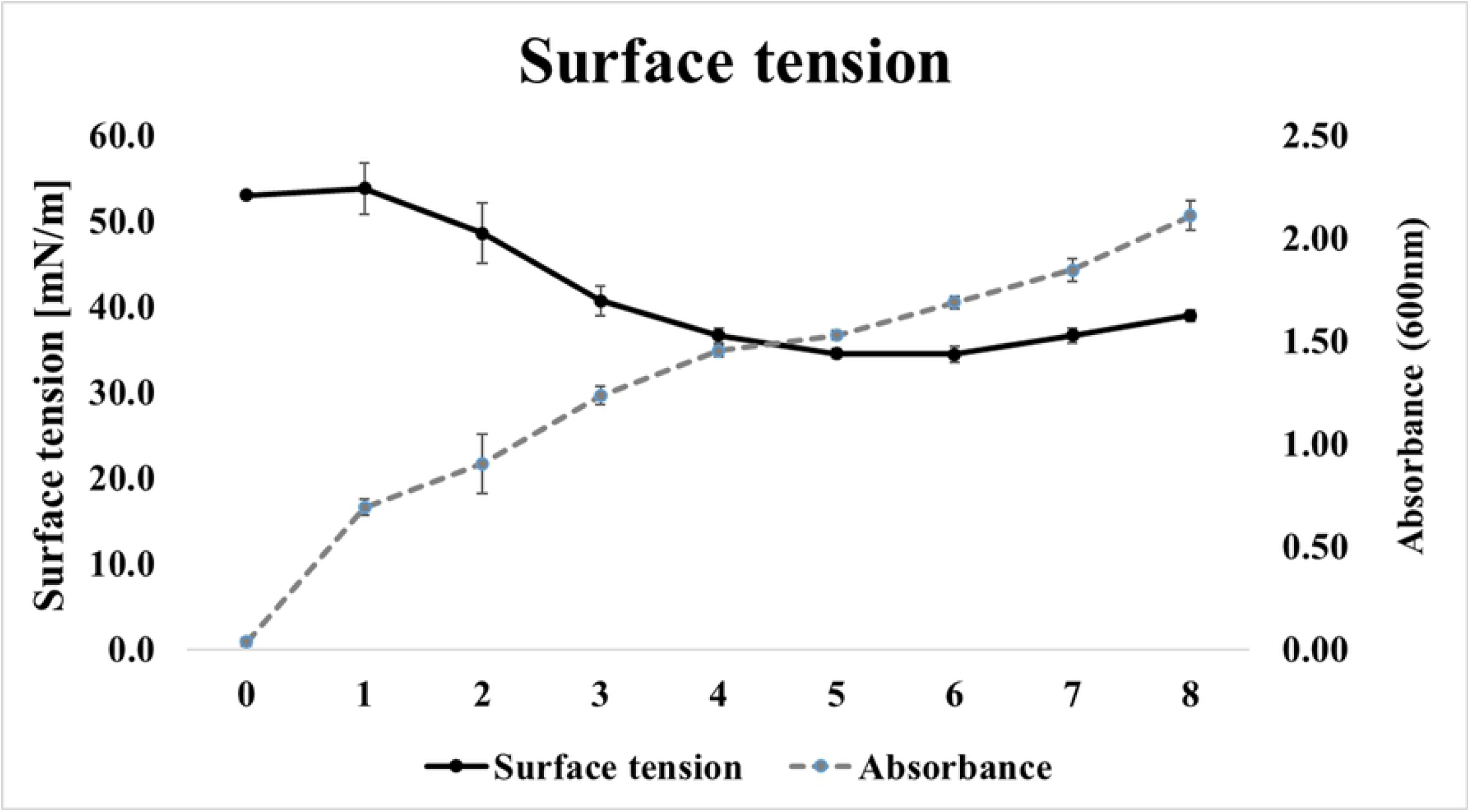
Time course of growth kinetics and surface tension in culture medium during cultivation of strain JAF-11.

The critical micelle concentration (CMC) is defined as the concentration of surfactant required to start the micelles formation. It is determined by plotting the surface tension measured according to the concentration of biosurfactant and identifying the point at which the surface tension of the biosurfactant no longer decreases dramatically. As a results of measuring surface tension of the water with the crude biosurfactants isolated from culture medium, the values were from 72.23 mN/m to 32.80 mN/m and the minimum surface tension value was 32.80 mN/m. In particular, the value of CMC was 24 mg/L that the concentration of the biosurfactant obtained from slope of the curve abruptly changed as shown in Figure 4.

**Fig 4.**
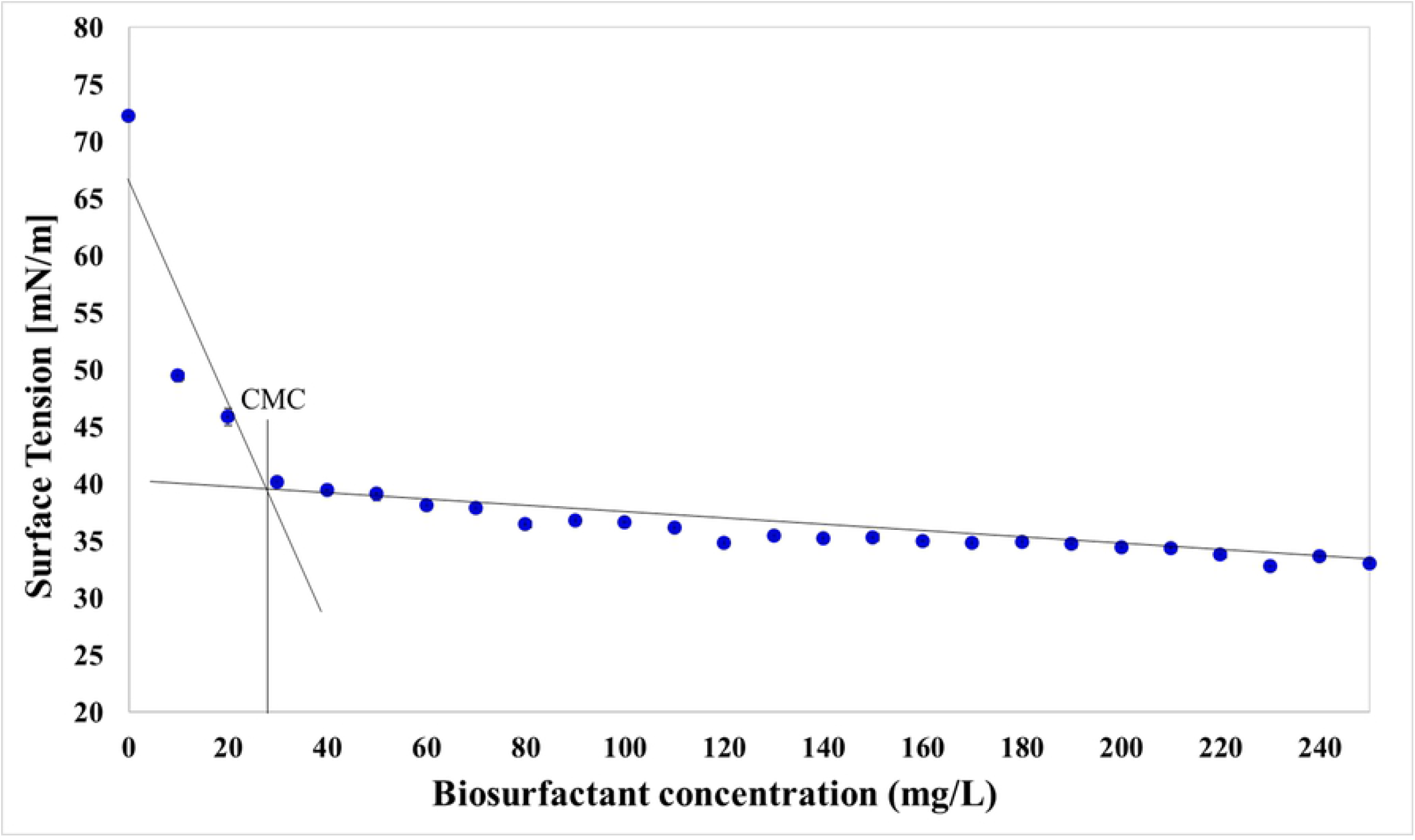
Determination of critical micelle concentration of producing crude biosurfactant from strain JAF-11.

### Chemical structure of the isolated compound

Chemical structure of the biosurfactant isolated was determined by mass and NMR measurements. The molecular weight of 502 was determined by the FAB-MS measurement, which showed a quasi-molecular ion peak at *m/z* 503 [M+H]^+^ (Fig 5). The molecular formula, C_26_H_46_O_9_, was determined by the HR-FAB-MS providing a molecular ion peak at *m/z* 503.3243 [M+H]^+^ (calcd. for C_26_H_47_O_9_, 503.3220), indicating four degrees of unsaturation. The ^1^H NMR spectrum of **1** (Table 2) showed signals due to six oxygenated methines at *δ*_H_ 5.50 (t, *J* = 2.7 Hz, H-2), 5.28 (t, *J* = 10.0 Hz, H-4), 4.93 (dd, *J* = 10.0, 2.7 Hz, H-3), 3.66 (t, *J* = 9.5 Hz, H-6), 3.65 (dd, *J* = 10.0, 2.7 Hz, H-1), and 3.45 (t, *J* = 9.0 Hz, H-5). It also showed signals attributable to 14 methylenes at *δ*_H_ 2.45 (m, H-2’)/2.42 (m, H-2’), 2.36 (m, H-2”)/2.31 (m, H-2”), 2.18 (m, H-2”’), 1.68 (m, H-3’), 1.59 (m, H-3”), 1.55 (m, H-3”’), and 1.35-1.25 (overlapped) and three methyls at *δ*_H_ 0.90 (overlapped). The ^13^C NMR spectrum (Table 2) in combination with HMQC spectrum displayed signals due to three carbonyl carbons at *δ*_C_ 175.0 (C-1’), 174.6 (C-1”), and 174.2 (C-1”’), six oxygenated methine carbons at *δ*_C_ 74.6 (C-6), 74.2 (C-5), 73.2 (C-4), 72.6 (C-2), 71.7 (C-3), and 70.9 (C-1), 14 methylene carbons at *δ*_C_ 23.4–35.2, and three methyl carbons at *δ*_C_ 14.4 (C-6”’) and 14.2 (C-8’ and C-6”). The ^1^H-^1^H COSY correlations among six oxygenated methine protons established the presence of an inositol moiety. Inositol moiety was identified as a *myo*-inositol by the proton coupling constant. Except for an equatorial proton at *δ*_H_ 5.50 (H-2) with coupling constant of 2.7 Hz, other protons occupied an axial position based on their proton coupling constants. The ^1^H-^1^H COSY spectrum also established six partial structures in three acyl chains, as shown in Fig 6B. Chemical structure was unambiguously determined by the HMBC spectrum, which exhibited the long-range correlations from two methyl protons at *δ*_H_ 0.90 (H-8” and H-8”’) to two methylene carbons at *δ*_C_ 32.4 (C-4” and C-4”’), from the methylene proton at *δ*_H_ 1.59 to the carbonyl carbon at *δ*_C_ 174.6 and the methylene carbons at *δ*_C_ 32.4 and 23.4, and the methylene proton at *δ*_H_ 1.55 to the carbonyl carbon at *δ*_C_ 174.2 and the methylene carbons at *δ*_C_ 32.4 and 23.4, implying the presence of two hexanoyl moieties. Other HMBC correlations from the methylene protons at *δ*_H_ 2.45/2.42 and 1.68 to the carbons at *δ*_C_ 175.0 and 30.2 and from the methyl protons at *δ* 0.90 to the carbon at *δ*_C_ 32.9, and 14 methylenes in the ^1^H and ^13^C NMR spectra indicated the presence of an octanoyl moiety. Finally, the long-range correlations from the oxygenated methines at *δ*_H_ 5.50, 4.93, and 5.28 to the carbonyl carbons at *δ*_C_ 175.0, 174.2, and 174.6, respectively, revealed that C-2, C-3, and C-4 in inositol moiety were acylated with one octanoyl and two hexanoyl groups, as shown in Fig 6B. Taken together, the structure of compound **1** was determined to be a new myo-inositol derivative and was named JAF-11. This compound was very similar to pullusurfactan E isolated from *Aureobasidium pullulans* strain A11211-4-57 from fleabane flower, *Erigeron annus* (L.) pers. [26], except for the presence of octanoyl moiety instead of hexanoyl moiety in pullusurfactan E (Fig 6A). Although the chemical structure is similar to pullusurfactan E, this study reports the isolation of a new biosurfactant from the novel yeast strain JAF-11 for the first time.

**Fig 5.**
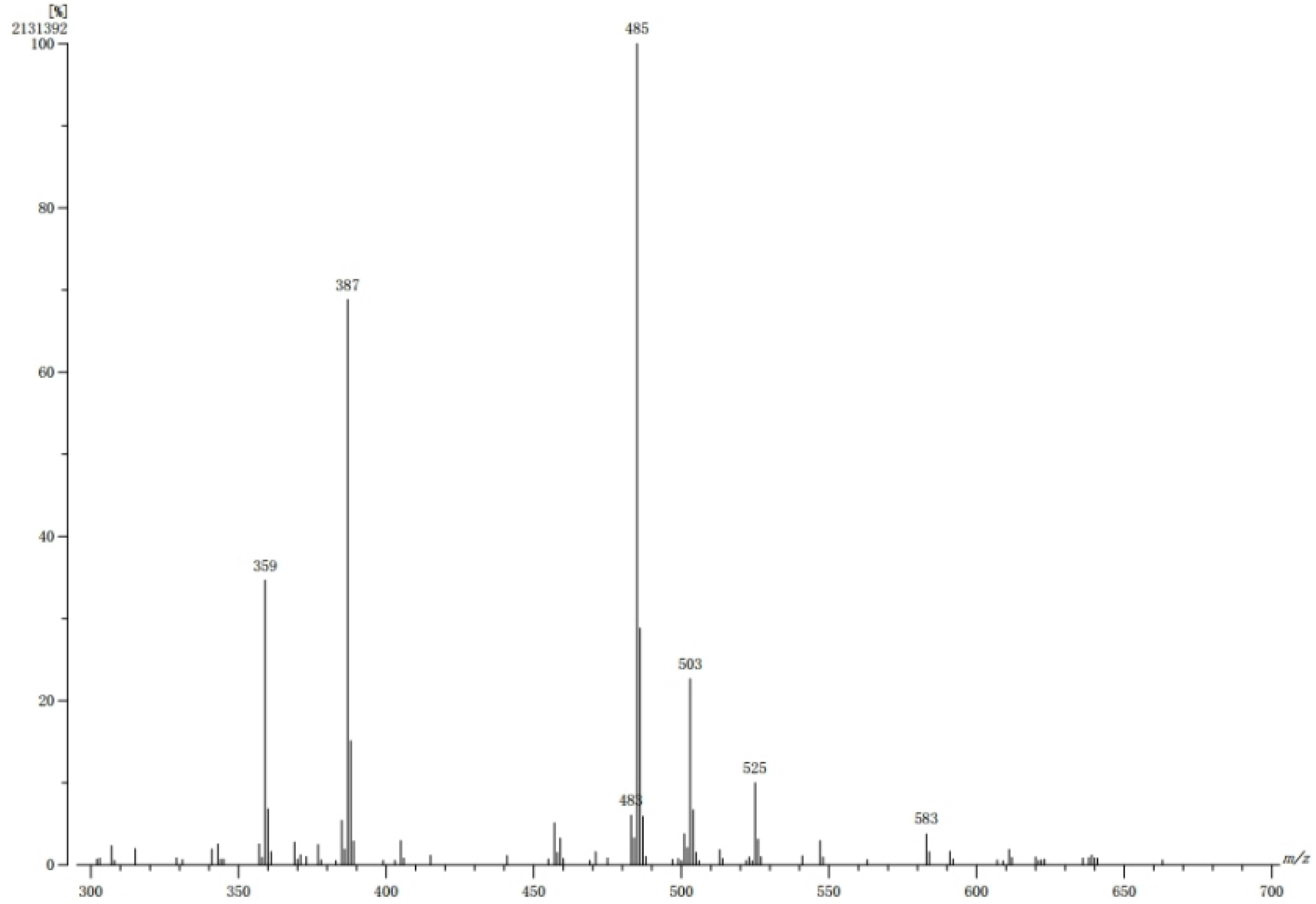
Fast atom bombardment mass spectrum of purified biosurfactant in the positive ion mode.

**Table 2.** ^1^H and ^13^C NMR spectral data of new biosurfactant in CD_3_OD.

| No. | $\delta_{\text{C}}$ | $\delta_{\text{H}}$ |
| --- | --- | --- |
| 1 | 70.9 | 3.65 (1H, dd, $J = 10.0, 2.7$)<sup>a</sup> |
| 2 | 72.6 | 5.50 (1H, t, $J = 2.7$) |
| 3 | 71.7 | 4.93 (1H, dd, $J = 10.0, 2.7$) |
| 4 | 73.2 | 5.28 (1H, t, $J = 10.0$) |
| 5 | 74.2 | 3.45 (1H, t, $J = 9.0$) |
| 6 | 74.6 | 3.66 (1H, t, $J = 9.5$) |
| 1' | 175.0 |  |
| 2' | 35.2 | 2.42 (1H, m), 2.45 (1H, m) |
| 3' | 26.3 | 1.68 (2H, m) |
| 4' | 30.2 | 1.38 (2H, m) |
| 5' | 30.2 | 1.25-1.35 (2H, m) |
| 6' | 32.9 | 1.25-1.35 (2H, m) |
| 7' | 23.7 | 1.25-1.35 (2H, m) |
| 8' | 14.2 | 0.90 (3H, overlapped) |
| 1'' | 174.6 |  |
| 2'' | 35.1 | 2.31 (1H, m), 2.36 (1H, m) |
| 3'' | 25.7 | 1.59 (2H, m) |
| 4'' | 32.4 | 1.32 (2H, m) |
| 5'' | 23.4 | 1.25-1.35 (2H, m) |
| 6'' | 14.2 | 0.90 (3H, overlapped) |
| 1''' | 174.2 |  |
| 2''' | 35.0 | 2.18 (2H, m) |
| 3''' | 25.4 | 1.55 (2H, m) |
| 4''' | 32.4 | 1.28 (2H, m) |
| 5''' | 23.4 | 1.25-1.35 (2H, m) |
| 6''' | 14.4 | 0.90 (3H, overlapped) |

<sup>a</sup> Proton resonance integral, multiplicity, and coupling constant ( $J$ =Hz) in parentheses

**Fig 6.**
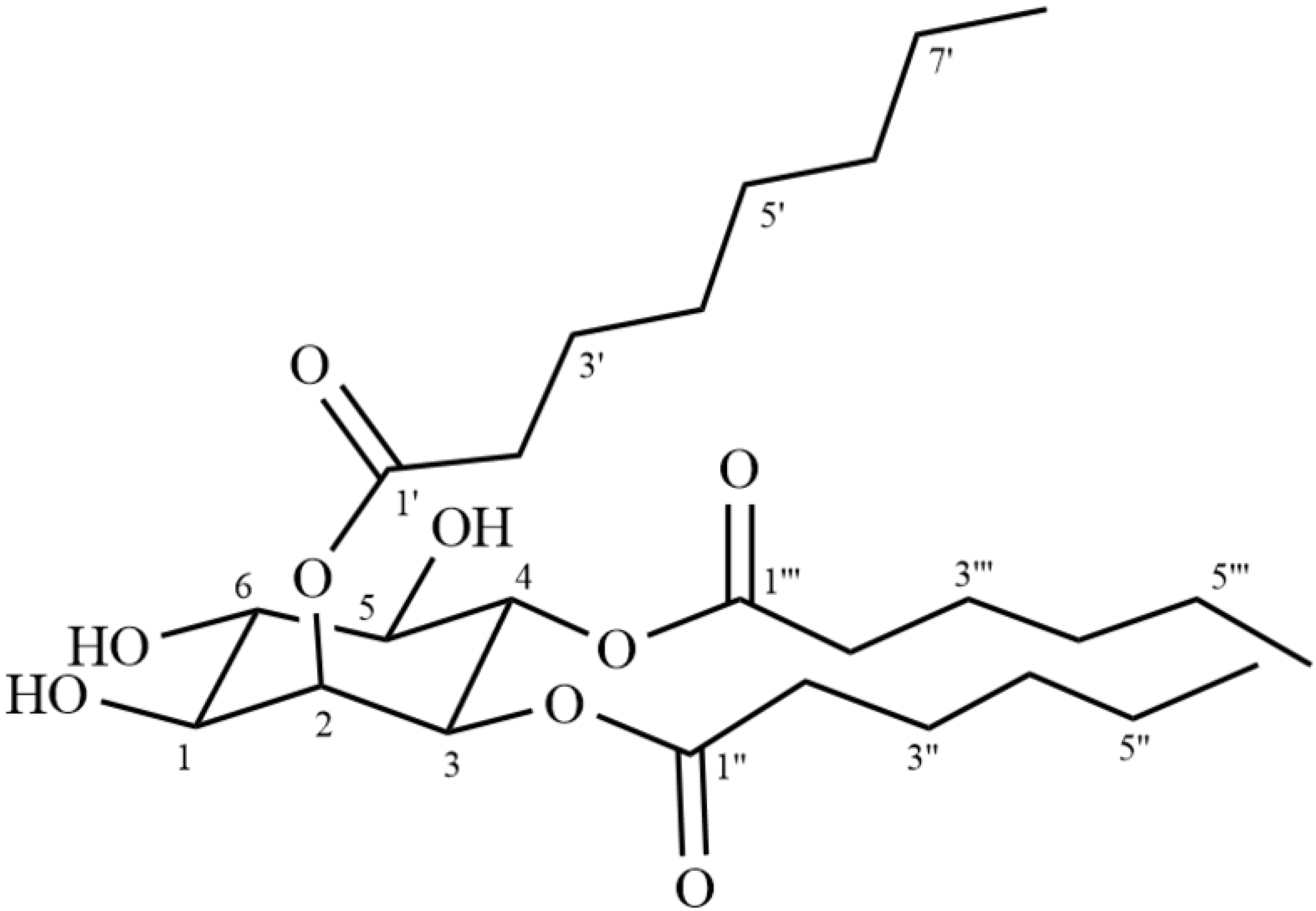
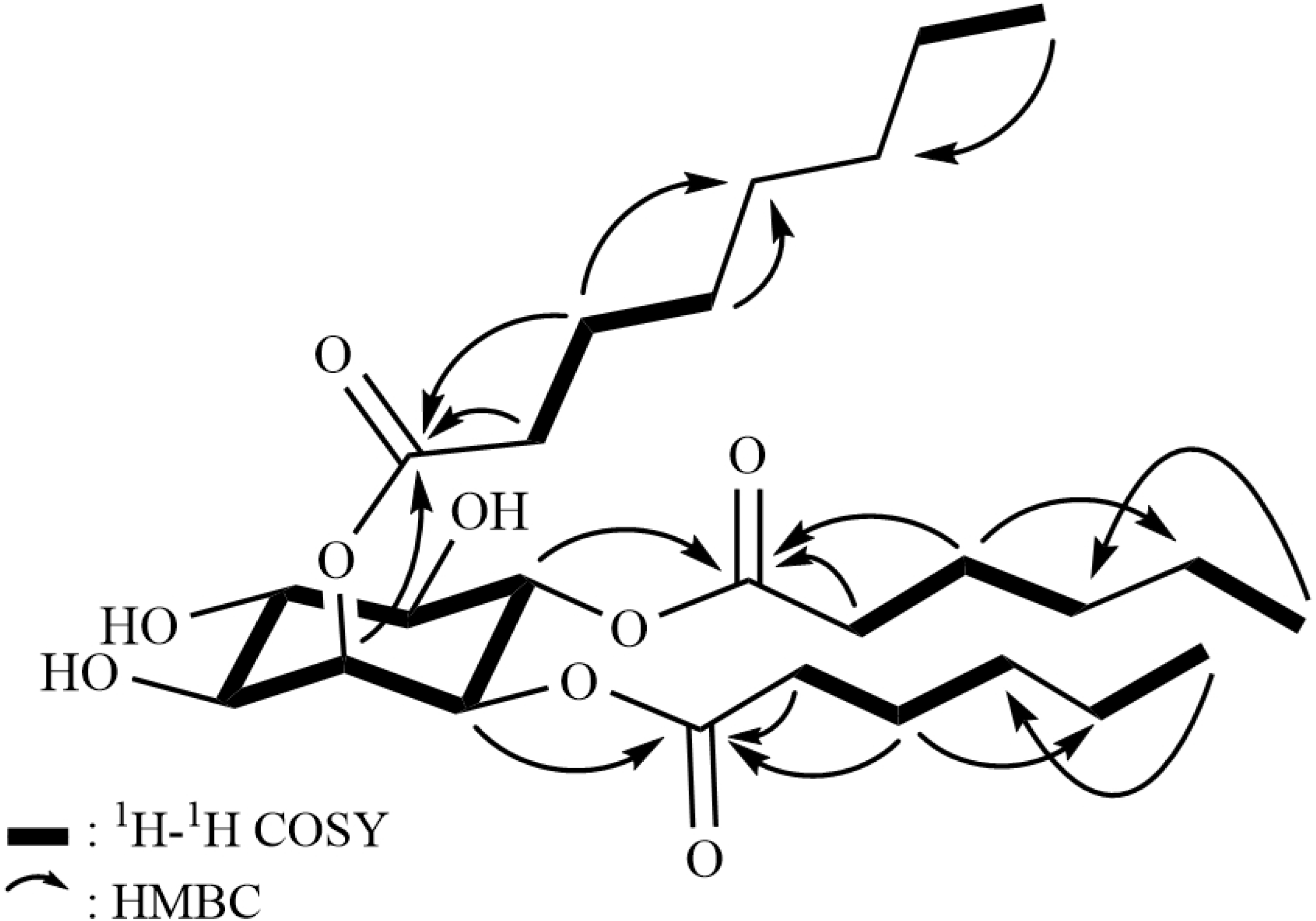
(A) Chemical structure of novel biosurfactant and (B) two-dimensional NMR correlations of novel biosurfactant.

## Acknowledgments

Authors thank Ms. Ji-Young Oh, Center for University-wide Research Facilities (CURF) at Jeonbuk National University, for performing NMR measurements.

## Supporting information

**S1 Fig. Screening of biosurfactant producing yeast by drop collapse method**. Distilled water and fresh culture broth were used as a control.

**S2 Fig. ^1^H NMR spectrum of the purified biosurfactant**

**S3 Fig. ^13^C NMR spectrum of the purified biosurfactant**

**S4 Fig. HMQC spectrum of the purified biosurfactant**

**S5 Fig. ^1^H-^1^H COSY spectrum of the purified biosurfactant**

**S6 Fig. HMBC spectrum of the purified biosurfactant**

